# Multi-drug interaction target-controlled infusion from simultaneous optimization of pharmacometric models

**DOI:** 10.1101/2024.03.24.586464

**Authors:** Pablo Martinez-Vazquez, Ana Abad-Torrent

## Abstract

**Background:** Accurately controlling drug delivery is crucial for safe anesthesia. Target-controlled infusion (TCI) systems use pharmacokinetic and pharmacodynamic (PK/PD) models to administer intravenous agents to reach target concentrations. However, TCI’s operation is restricted to a single-drug and a linear PK/PD model, not accounting for drug interactions. Dose-response interaction (DRI) models quantify such interactions by representing shared effects as a function of agents’ concentrations. For example, the co-administration of an analgesic and a hypnotic with TCI leads to an uncontrolled synergy.

**Methods:** We introduce a new administering methodology for multi-drug infusions, interaction target-controlled infusion (iTCI), that combines the PK/PD models of the co-administered drugs and their interactions into a single optimal non-linear dynamic control problem with terminal constraints.

**Results:** Incorporating DRI and PK/PD models allows novel administration procedures. Simulations of iTCI in different clinical scenarios under propofol and remifentanil co-administrations are presented. These show that: (1) iTCI requires lower administered volumes than TCI to reach simultaneously the same target concentrations. (2) It offers optimal interdependent administrations that address not only concentration targets but also effect targets. (3) iTCI comes with additional constraints on the administration, including controlled titrations along iso-effect conditions (isoboles) or (5) directly limiting plasma concentration levels. (6) Unlike TCI, iTCI can include different exerted effects (ke0) per drug, particularly relevant for opioids.

**Conclusion:** The iTCI is a versatile multi-drug infusion paradigm where effects and interactions play a relevant role - providing better delivery profiles than current TCI while opening the door for using non-linear PK/PD descriptions in anesthesia.

## 1 INTRODUCTION

Traditionally, intravenous (IV) have been administered according to standard dosing guidelines, either as boluses or continuous infusions, until the advent of the target-controlled infusion (TCI) systems in the early 1990s, which tailored drug administrations to patient characteristics (age, sex, weight, and height), allowing to control continuously the accumulation of the drug in the body tissue from automatic optimal drug delivery actions.^1,2^ TCI use has proven to be as safe and effective as manually infusions, but with rapid and accurate drug titrations to achieve desired effects under quick surgical changes, and maintaining steady concentrations during periods of constant stimulation.^3–6^ TCI-guided anesthesia resulted in better vital signs and lesser post-operative nausea and vomiting episodes.^3,6,7^

A TCI system is a computer-guided pump based on a pharmacokinetic/pharmacodynamic (PK/PD) model dosing an IV agent to achieve a desired target concentration in a tissue or site of interest in minimal time. In general anaesthesia (GA), the two drug concentrations accounted for in a PK/PD analysis are the plasma concentration (Cp) and the effect-site concentration (Ce). Cp is the concentration of a drug in the blood plasma after considering various pharmacokinetics factors such as absorption, distribution, metabolism, and elimination. The Ce represents the drug concentration at its action site, evaluated by the pharmacodynamics. Depending on the clinical context, once the anesthesiologist establishes a desired concentration target, either in Cp or Ce, the TCI automatically calculates the upcoming infusion administration actions to rapidly achieve the selected target based on the continuously updated estimations of current and previous concentrations in the body sites from a PK/PD drug model tailored to the patient’s characteristics.^2,5^

TCI systems deliver in an automatized way: a single drug. The control operates independently of other co-administered agents that may interact with the controlled anesthetic. Besides, TCI does not directly include effects that the drug exerts in the control strategy, making TCI a sub-optimal delivery solution missing important clinical aspects.

Anesthesia is the practice of applied drug interactions.^2,8^ Drugs are rarely administered alone, and different drugs interact with each other in such a way that rarely does a drug, when administered in the presence of other drugs, behave as if it were administered alone. Therefore, when multiple interacting drugs are administered simultaneously, it is advisable to integrate their respective body site concentrations, infusion rates, and interactions into a common control. This approach results in better dosing outcomes for all drugs compared to using separate TCI controls for each one.

Under this principle, we introduce a novel multi-drug administration method called interaction target-controlled infusion (iTCI). Unlike TCI, iTCI provides several advantages: (1) achieving target levels for all drugs simultaneously, reducing overall drug usage; (2) faster target achievements; (3) allowing constrained transitions between dosing regimes under iso-effect (isobole) conditions, and (4) direct concentration constraints. Among others, these benefits provide practitioners with enhanced control over drug dosages, towards improving anesthesia safety.

## 2 METHODS

### 2.1 ALGORITHM DESCRIPTION

The article targets anesthesiologists, outlining the clinical benefits of this new drug control framework. The iTCI description follows a brief basic overview of pharmacometric models and TCI concepts needed to aid understanding. Examples of iTCI’s application in relevant clinical scenarios are provided, emphasizing each of its main advantages over TCI.

### 2.2 The PK/PD model. The basis of TCI

TCI relies on empirical PK/PD drug models that models describe how drugs move through the body and their effects..^2,5^ The PK component depicts the time course of drug distribution, absorption, and elimination in the body. It is typically represented by a three-compartment model with a central compartment of volume (V1), where the drug is injected, connected to two peripheral compartments (V2 and V3), representing fast and slow redistribution, respectively, fig. 1.A.

**Figure 1:**
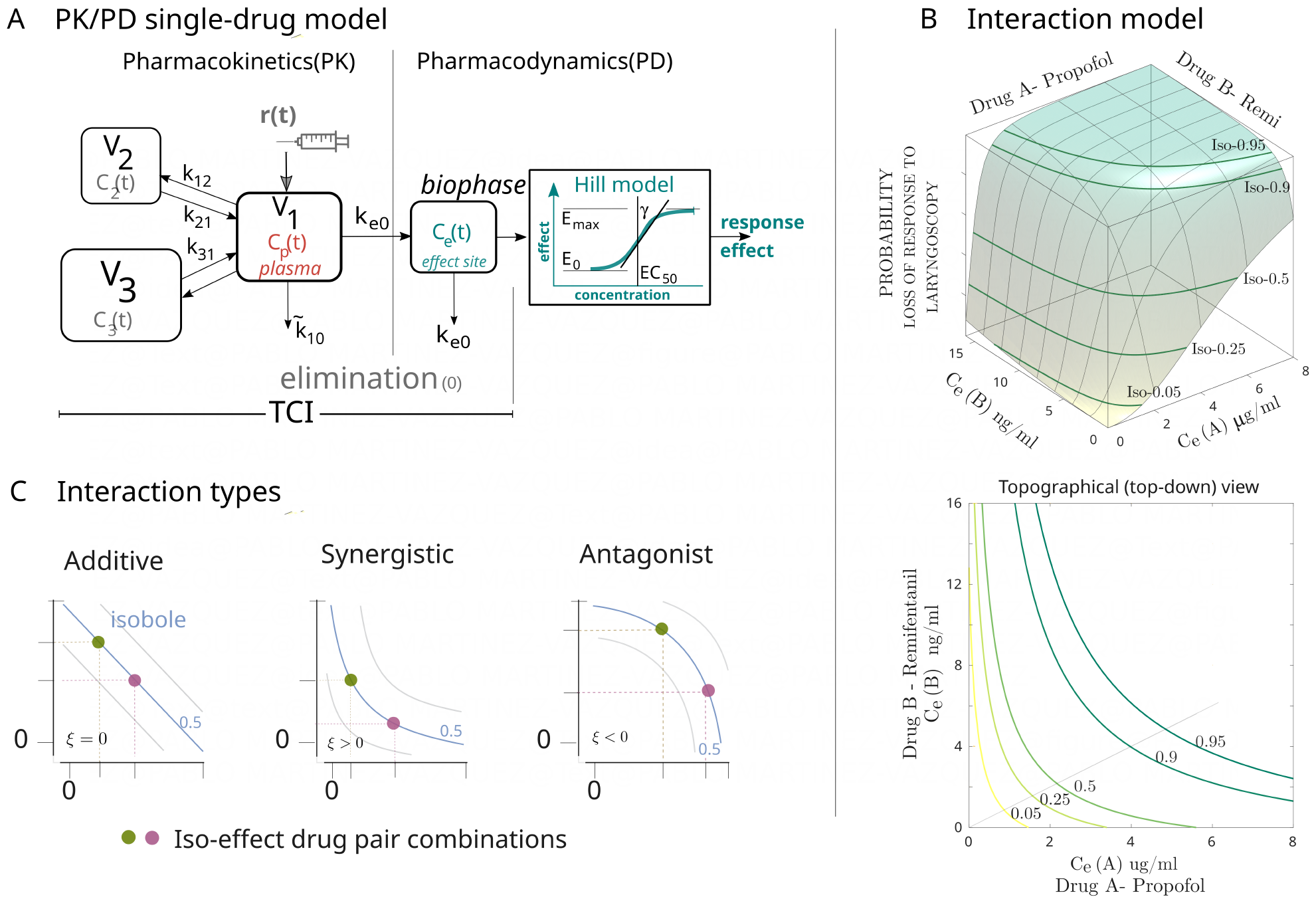
(A) The PK/PD model is a dynamic description of the pharmacokinetics and pharmacodynamics of the drug. The PK component consists of three compartments or volumes. A central compartment (V1), into which the drug is injected, is connected to two peripheral compartments that represent the fast (V2) and slow (V3) distributions of the drug in the body. Plasma concentration Cp(t) is a function of the infusion rate (r(t)), the drug concentrations and distributions across the peripheral compartments, and its elimination. The mass transfer rates, *k*_*ij*_, characterize the inter-compartment drug distributions. Drug elimination is modeled as a diffusion to an external body compartment (0). The PD part describes the evolution of the effects produced by the drug. It is a theoretical compartment representing the drug concentration at the biophase (Ce), and a nonlinear function that translates the Ce into an effect, commonly formulated with the Hill equation. The virtual Ce compartment, connected to V1, accounts for the delay T_1/2_ between Cp and drug effect. The concentration-effect relationship is not used in TCI but in ke0 estimation. (B) Interaction models describe how two or more drugs work together. The dose-response interaction (DRI) is a generalization of the single drug concentration-response model to several interacting agents, fundamentally, drug-drug interactions. It describes the relationship between two Ce on a common effect or response. The three-dimensional DRI example corresponds to the propofol-remifentanil interaction of loss of response to laryngoscopy, with its topographical view below. The pairs of iso-effect concentrations are depicted by level lines (isoboles) of constant probability of loss of response to laryngoscopy. (C) Types of drug-drug interactions and isobole shapes at 50% of response level. If the overall effect of the drugs is the sum of their individual effects, they have an additive interaction. If the overall effect is greater than the sum of their individual effects, they interact synergistically. Conversely, if the overall effect is less than the sum of their individual effects, they interact antagonistically.

The drug concentration in the central volume (C_1_(t)or C_p_(t)) depends on infusion rate (r(t)), distribution to peripheral compartments, and elimination to an external compartment (0). These factors are modeled by linear differential equations where C_2_(t) and C_3_(t) represent the concentrations in the peripheral compartments, eq. (1). Constant k_ij_ ([min^*−*1^]) denotes the mass transfer rate from compartment i to compartment j, and 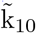 accounts for the drug elimination rate. The injected drug concentration in mass per volume units is denoted by *I*.

Distinct PK models or fittings for different drugs share this mathematical approach, differing in the volumes and k_ij_ estimates expressed as functions of patient characteristics.^2^ The TCI execution adapts to the model parameters’ estimates. Table A-1 shows two PK models for propofol and remifentanil, respectively.

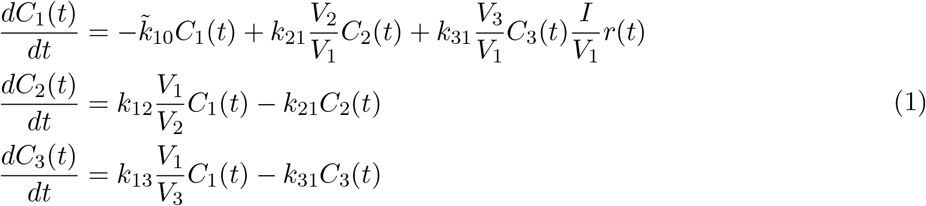

The PD aspect examines the relationship between Cp and an exerted effect. As the plasma is not the drug’s site of action, a delay between Cp and effect is observed. This delay is represented by a theoretical compartment connected to the central volume, which represents the drug concentration in the biophase, Ce. To describe the equilibrium between these two compartments, a first-order inter-compartmental transfer rate (ke0), eq. (2a) is used. This rate can be expressed as the equilibrium half-time delay T_1*/*2_(ke0) = ln(2)*/*ke0, accounting for the time Ce needs to reach 50% of Cp when Cp remains constant.

The impact of Ce on a specific effect (E), or concentration-effect relationship, is modeled by a nonlinear function, typically the Hill model, fig. 1. An Emax-sigmoidal function, eq. (2b), where E_0_ represents the effect under drug absence, E_max_ is the highest effect, EC_50_ denotes the concentration associated with half the difference between E_max_ and E_0_, and *γ* indicates the steepness of the relationship. Any effect is measured from vital signs or patient responses such as heart rate, blood pressure, EEG or algometry, and a metric. Unlike PK, which relies solely on blood-sampled data, PD modeling and its interpretation depend on the chosen response and metric.^2,9^

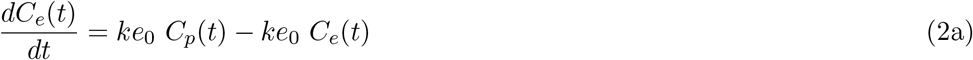

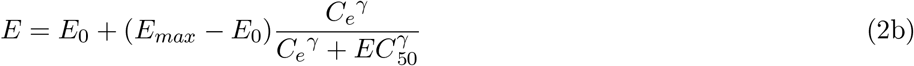

### 2.3 TCI as an optimal control problem

TCI is a computer-guide pump that executes an infusion profile to bring current drug concentrations, particularly Cp and Ce, to some desired target values in a minimum time. The TCI action can be formulated as the solution of a linear dynamic optimization problem with terminal constraints, eq. (3), where the PK/PD model, eqs. (1) and (2a), was compactly rewriting into a matrix form 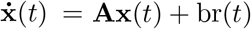. The state vector, **x**(*t*) = [C_1_(t), C_2_(t), C_3_(t), C_e_(t)]^*T*^, contains the drug compartmental concentrations at time t, and the matrix **A** and vector b are formed with the transfer mass rates (k_ij_) and compartment volumes (V_i_).

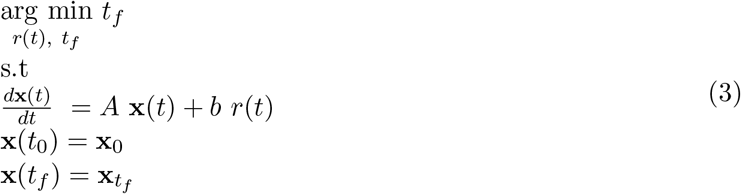

The TCI action is the infusion rate (*r*(*t*)) over a time *t* (*t*_0_ *≤ t ≤ t*_*f*_), required to bring the drug concentrations from a given initial state (**x**(*t*_0_) = **x**_0_) to final desired values, 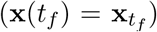, in a minimum time (*t*_*f*_), subject to (s.t) a given PK/PD model. A minimization (arg min) of target reaching time *t*_*f*_ subject to all given conditions: initial state, final state or target, and the PK/PD model. TCI offers two modes of operation: Cp-target and Ce-target mode. Both modes are obtained by changing the end conditions of 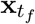in eq. (3).

Closed-form solutions to this optimization problem of end-point constraints were published,^10–13^ with Shafer and Gregg’s solution becoming the basis of today’s modern TCI pumps.^1,11^

### 2.4 Drug interactions and modeling

Although each anesthetic primarily targets a specific major component of anesthesia (e.g., hypnosis, analgesia, muscle relaxation), its influence is rarely limited to just one component. For instance, administering an opioid like remifentanil with a hypnotic like propofol produces deeper hypnosis and analgesia than using them separately.^8,14–17^ Therefore, it is important to consider drug interactions in anesthesia administration to achieve better control of their effects on the body

Dose-response interaction (DRI) models are quantitative descriptions of net clinical endpoints that result from administering multiple drugs based on their Ce concentrations. While there is flexibility in choosing and quantifying different effects, in GA, the foremost effects of interest are evaluating the patient’s response to particular external stimuli. These responses may include reactions to verbal, tactile, pressure algometry, or tetanic stimuli, as well as changes in hemodynamics, electrical brain activity, and reactions to laryngoscopy tracheal intubation, LMA insertion, or esophageal instrumentation.^9,15–20^

DRI models are often created using either Greco, eq. (4), or Minto functions, both of which are generalizations of the single-drug Emax-sigmoidal function, eq. (2b).^20,21^ Particularly, the drug-drug DRI model is a three-dimensional surface in which the x- and y-axes represent the two Ce concentrations, and the z-axis represents the patient’s response.

The drug-drug Greco model measures the combined effect of two drugs - A and B, *E*_*AB*_. It is a function of the maximal effect, E_max_, along with the individual drug concentrations that produce 50%of the maximum effect, 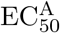 and 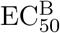, and the interaction strength *n*. The *ξ* characterizes the nature of the interaction. An *ξ* = 0 indicates additive interactions, while *ξ <* 0 and *ξ >* 0 mean antagonistic and synergistic interactions, respectively.

Fig. 1B shows the DRI model of *E*_*AB*_, defined as the probability of loss of response (LOR) to laryngoscopy, for propofol and remifentanil given by Kern et al..^15^ The interaction between propofol and remifentanil is clearly synergistic. Particularly, relevant in clinic are the iso-effect curves (isoboles) corresponding to the continuum of drug pair concentrations leading to the same net effect or response. Isoboles of clinical interest during GA are just above, but not far beyond 90-95%.

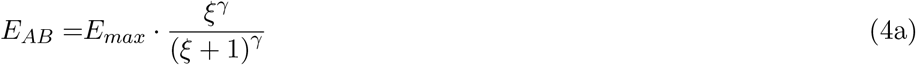

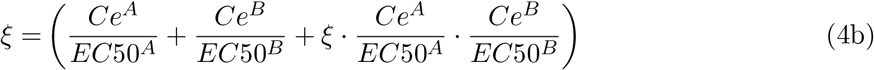

DRI models are powerful representations of drug interactions composed of the full range of isoboles of all drug combinations.^15,21,22^ They will enable the connection between distinct single-drug PK/PD models in the iTCI control. Although iTCI is applicable for n-drugs, its presentation and simulations will deal only with two-drug infusions, as they are the current modeled and studied interactions.^8,20,21^

## 3 iTCI - ALGORITHM DESCRIPTION

Unlike TCI, in which each drug administration profile is computed independently of the other administered anesthetics, in the iTCI, the administration profiles for all co-administered drugs are calculated by integrating their respective PK/PD models and interactions into a global control solution; a generalization of the TCI optimizations that provide better infusion regimes. Fig. 2 shows a schematic comparison between the TCI and iTCI for two co-administered interacting agents, A and B.

**Figure 2:**
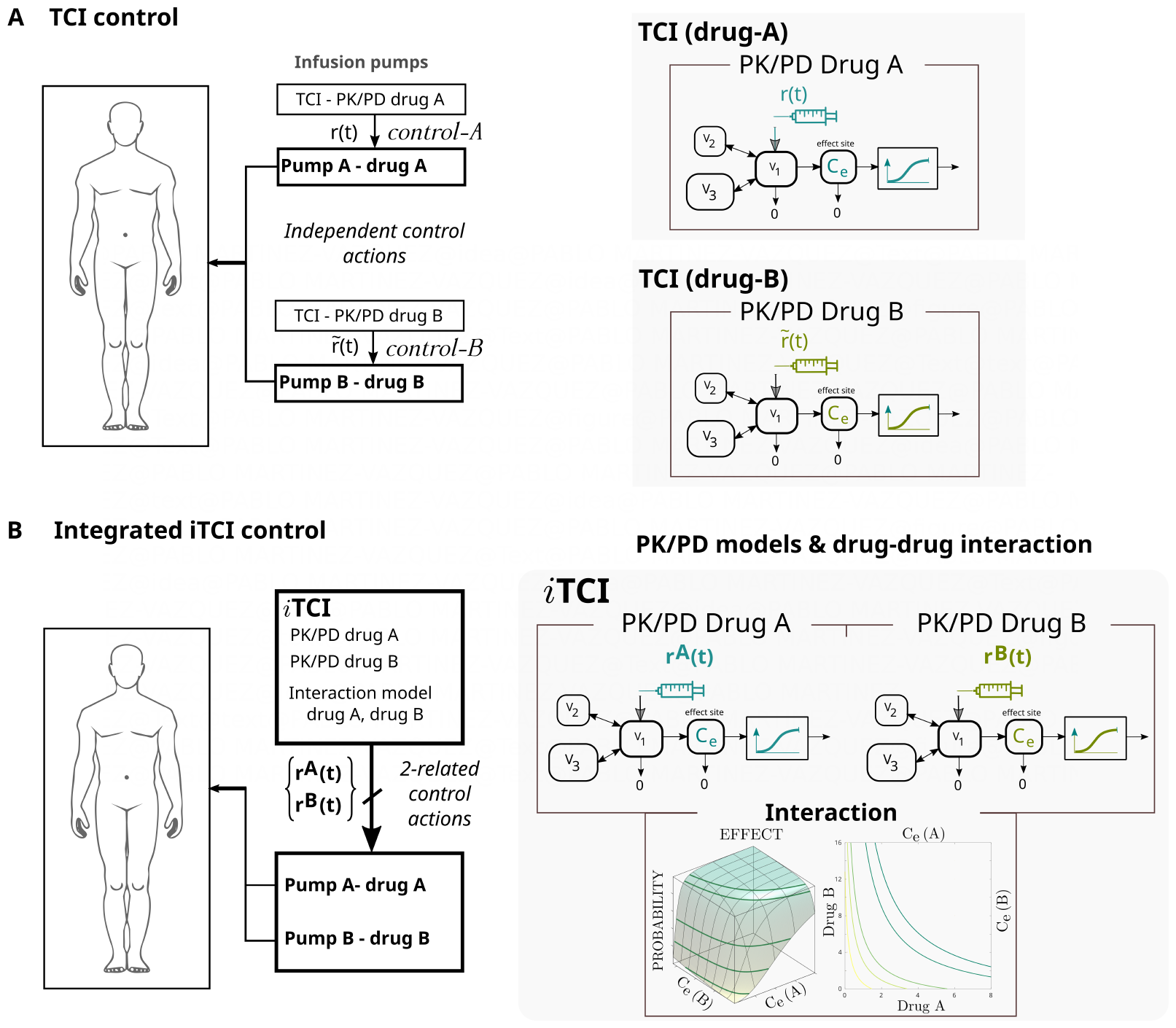
Schematic comparison between TCI and the novel framework for the case of two drug concurrent administrations. (A) The current TCI system doses the two anesthetics with two independently functioning TCI pumps. Given the desired target concentrations, in C_p_ or C_e_, each drug is administered with an optimal control profile calculated regardless of the dose of the other interacting co-administered drug. (B) The iTCI integrates the delivery control of the 2-drugs into a single controller, which calculates the optimal delivery profiles for each pump, {*r*^*A*^(*t*), *r*^*B*^(*t*)}, concurrently from two of the PK/PD models and their drug-drug interaction. Therefore, each drug delivery profile depends not only on its PK/PD properties but also on the other drug delivery profile.

The linear dynamic optimization problem statement of TCI, eq. (3), can be generalized into a non-linear dynamic optimization problem with terminal constraints, with the general form of eq. (5), defining an augmented state-vector **x**(*t*) = [**x**^*A*^(*t*), **x**^*B*^(*t*), *E*_*AB*_(*t*))]^*T*^, consisting of each drug state-Vectors, 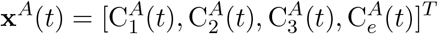 and 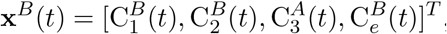, with the compartmental concentrations and their interaction effect-response, *E*_*AB*_(*t*).

The iTCI actions for two drugs, A and B, are defined, similarly to TCI as the optimal infusion profiles, **r**(*t*) = [*r*^*A*^(*t*), *r*^*B*^(*t*)]^*T*^, during time *t*_0_ *≤ t ≤ t*_*f*_, to bring Cp and Ce concentrations for each drug accounted in extended state-vector **x**(*t*), from given initial values, **x**(0) = **x**_0_, to final target values, 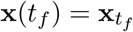, in a minimum time (*t*_*f*_), eq. (5). Optimal infusions, **r**(*t*), that fulfill (s.t-subject to)) the dynamical system 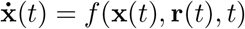, *t*) that includes the two PK/PD models 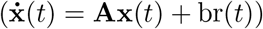, and their interactions under a given DRI model.

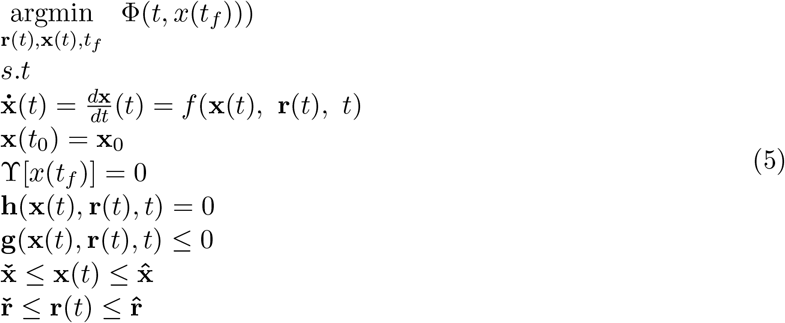

Given a patient initial state, **x**(0) = **x**_0_ (concentrations and effect), different control strategies will be obtained depending on the terminal target state-vector definition, 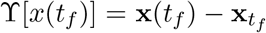, as well as additional extra equality and inequality constraints on the states or input controls. Conditions 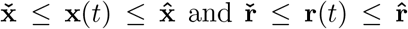 denote lower and upper limits for the states and infusion rates, respectively.

The differential-algebraic optimization problem, eq. (5), can be solved from a complete parametrization of controls, **r**(t), and states, **x**(t); converting this statement problem into an algebraic non-linear programming (NLP),^23–26^ that can be solved by a successive quadratic programming (SQP) solver.^27^ See A-2 for a detailed iTCI description.

## 4 RESULTS

### i TCI simulations

The iTCI function will be shown in clinically relevant scenarios during GA. Examples correspond to co-administration of propofol and remifentanil (designated as drugs A and B, respectively) to a 33 years old subject weighing 84 kg, with a height of 179 cm and LBM of 64.2 kg.^28^ Propofol and remifentanil concentrations were set to 10 mg ml^*−*1^ and 50 *μ*g ml^*−*1^, respectively, with maximum infusion rates of 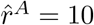 *ö*g min^*−*1^ and 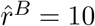 ng min^*−*1^.

The selected pharmacometric models correspond to (1) the PK/PD fittings for propofol and remifentanil by Schnider and Minto, respectively, and (2) the interaction model, DRI by Bouillon et al. for the LOR to laryngoscopy.^17,21,29,30^ Without loss of generality, iTCI operates likewise with different fittings.

The control examples include:

- Minimum reaching time to Ce-targets.
- Minimum time to reach a target effect.
- Transitions along isoboles.
- Direct constraints on concentrations.
- Multi-effect control.

### 4.1 Minimum reaching time to Ce-targets

The first example of iTCI shows a controlled infusion to a patient from a given state of concentrations, **x**(*t*_0_), to a new pair of desired Ce-target concentrations, 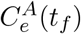 and 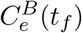. For simplicity, the initial state of concentrations was initialized to zero (**x**(*t*_0_) = 0), representing a complete absence of drugs in the body, simulating an anesthesia induction. The propofol and remifentanil Ce-targets were set to 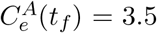 *ö*g ml^*−*1^ and 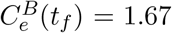 ng ml^*−*1^, respectively, corresponding to a 90% LOR to laryngoscope, according to the DRI model.

Fig. 3 displays the iTCI administration profile, showing the trajectory of propofol-remifentanil Ce concentrations on the interaction model from the initial state to the desired targets. The time evolutions of the four compartmental concentrations for the two agents are shown below, corresponding to the infusion rates given by the iTCI, along with the total administered volumes. The iTCI control ensures that both agents reach their targets simultaneously at time *t*_*f*_ = 2.1 min, taking into account both PK/PD properties.

**Figure 3:**
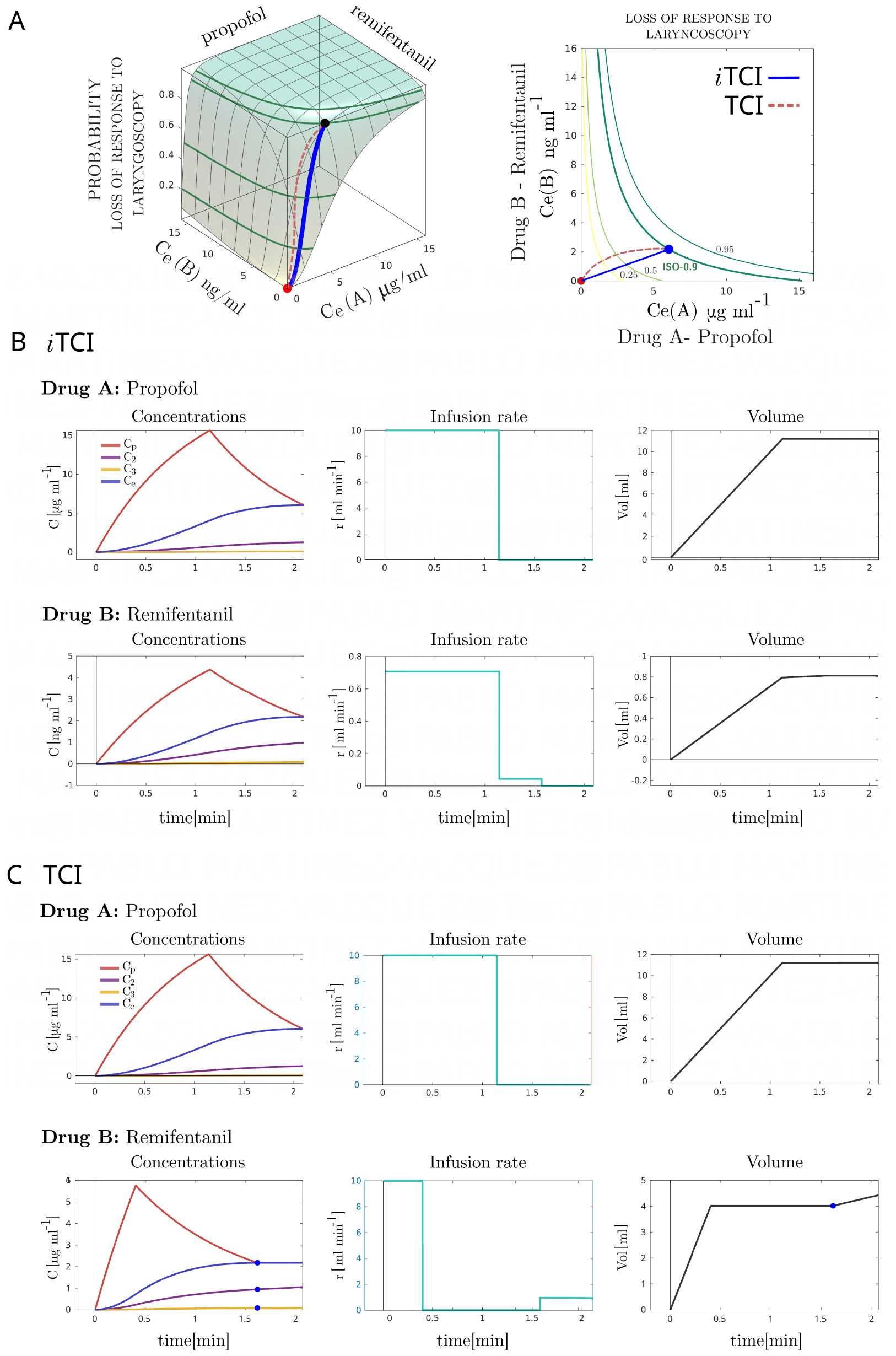
Minimum time to reach a pair of desired Ce-targets: Propofol target of 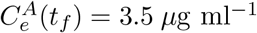 and remifentanil target of 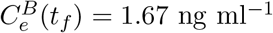, corresponding to a 90% LOR to the laryngoscope. (A) iTCI and TCI induction trajectories along the DRI model. 3-D representation and a top-down view. (B) iTCI optimal propofol-remifentanil profiles, including concentrations, infusion rates (control actions *r*(*t*)), and administered volumes over time. (C) Independent TCI optimal profiles for propofol and remifentanil. Concentrations, infusion rates, and administered volumes per drug over time.

In comparison, TCIs, with independent administration profiles per drug, lead to different reaching target times, Fig. 3C. Remifentanil reaches its target at 1.6 min, earlier than propofol, which is slower, requiring 2.1 min. However, achieving the desired pair of targets, aiming for the 90% LOR to the laryngoscope, can only be accomplished once the propofol goal is attained. Therefore, remifentanil must be kept constant.

The independent drug management of TCI directs to unneeded fast administration of the faster agent (remifentanil), resulting in unnecessary larger administered volumes. TCI delivers remifentanil in the largest bolus possible (max. infusion rate 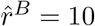 ng min^*−*1^) to achieve its target, however, in the iTCI strategy, since the other agent’s PK/PD characteristics and targets play a role, the profile for remifentanil is adapted to a much lower bolus, which translates in volume reductions and lower peak plasma levels. These differences in reaching target times and administered volumes become larger as the differences in drugs’ PK/PD characteristics increase. Fig. 3A shows the two TCI Ce trajectories (dash-red) in the Ce-space.

In iTCI, the drugs’ Ce-targets and infusion rates are always interrelated and sharing a common end-timing, *t*_*f*_. This condition is desirable because what matters most is the resulting patient state from all administered agents, rather than the effects of each agent on its own. Simultaneity and interrelation provide control strategies that lead to reduced consumption and plasma levels of the faster-acting agent(s).

### 4.2 Minimum time to reach a target effect

The iTCI’s ability to operate with general dynamic models of the form 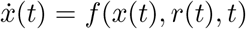, rather than being limited to linear ones, opens up the possibility to rethink the multi-agent control strategy beyond concentrations. Instead of focusing on Ce-targets (*Ce*^*A*^, *Ce*^*B*^), the algorithm can optimize infusions to attain a desired effect, without speechifying any Ce-targets, by including the interaction model, eq. (4), in the optimization problem and the desired *E*_*AB*_-target in the terminal condition, ϒ(*t*_*f*_).

Fig. 4 shows the iTCI induction with propofol and remifentanil, starting from **x**(*t*_0_) = 0, to rapidly achieve the targeted effect of *E*_*AB*_ = 0.9. The iTCI yields the fastest co-infusion (*t*_*f*_ = 1.72 *min*) to the desired effect, along with its corresponding ending Ce-pair (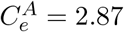 *ö*g ml^*−*1^,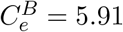 ng ml^*−*1^), obtained from all possible pairs within the 90% isobole of LOR to the laryngoscope.

**Figure 4:**
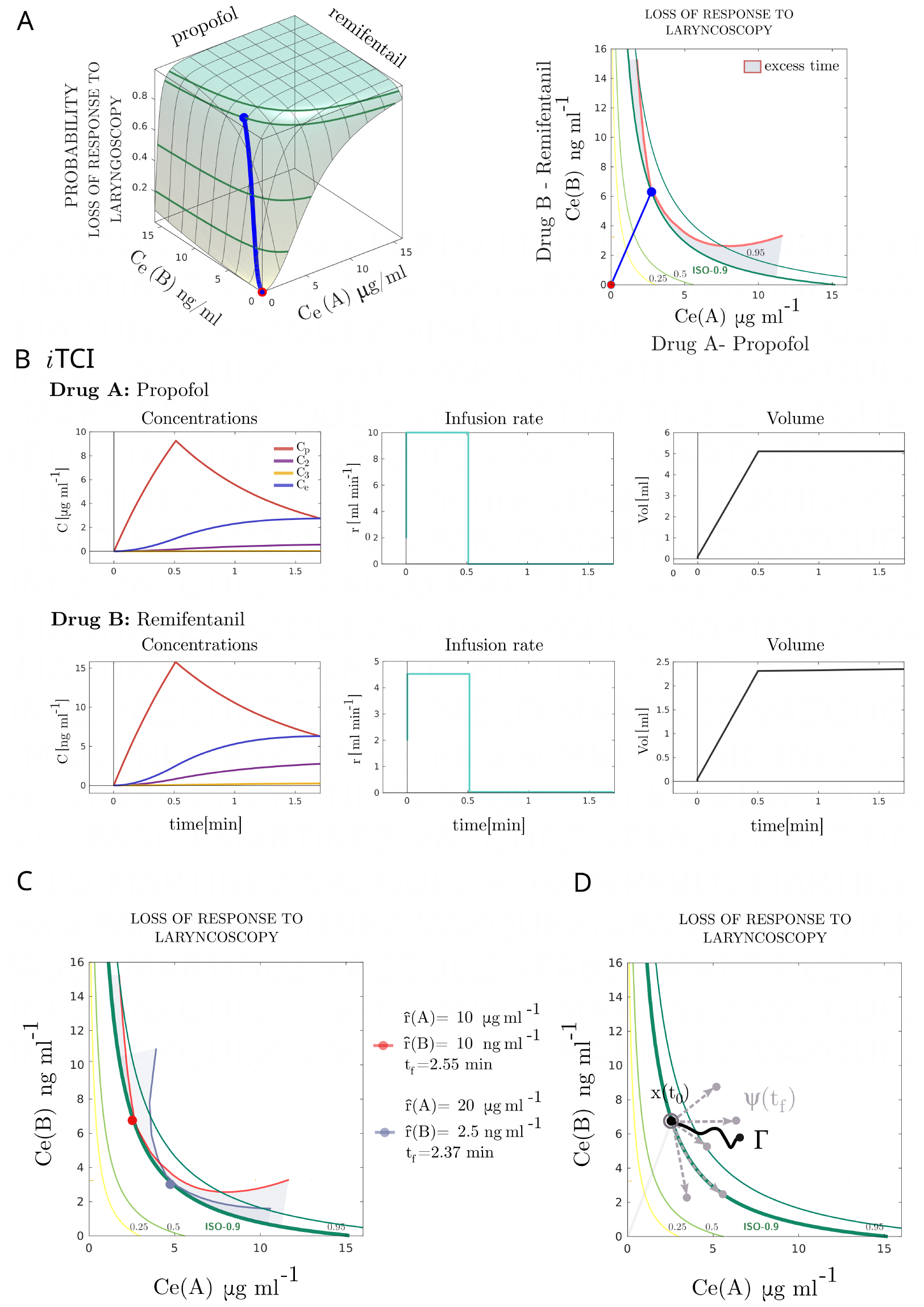
Minimum induction time to reach a desired target effect, *E*_*AB*_, represented by the isobole of the 90% of probability of loss of response to laryngoscopy. The iTCI algorithm identifies the optimal Ce-target pair from all the possible pairs within the isobole for the defined infusion conditions.(A) iTCI solution trajectory in the DRI model, shown in 3D and top-down views. The top-down view illustrates the excess time function for different Ce-target pairs on the target isobole. (B) iTCI infusion rates (*r*(*t*)) to the selected effect-target, displaying concentrations and administered volumes of drugs over time. Infusion rates were limited to maximum values of 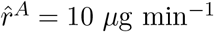 and 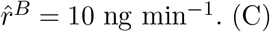. The minimum reaching time and location on the isobole is problem-dependent on infusion constraints. For instance, two different times and Ce-reaching pairs depend on the admissible maximum infusion rates. (D) Besides defining the multi-drug control problem from the initial and terminal desired states (*x*(*t*_0_) and Ψ(*t*_*f*_)), the iTCI framework allows for restricting the control strategy along a desired feasible trajectory (*Γ*) in the DRI model space starting from a given *x*(*t*_0_).

Compared with previous Ce-targeted iTCI, Fig. 3, the effect-targeted profile allows reaching the same effect in a much shorter time. Additionally, under the triad (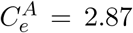 2.87 *ö*g ml^*−*1^, 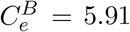 ng ml^*−*1^, *E*_*AB*_ = 0.9), the administered volumes are also lower. The effect-target iTCI is of particular interest since it finds the best end for desired effect than manually testing Ce-targets within an associated isobole. Effect-target broadens practitioners’ views to think not only in terms of concentration targets but also in terms of effects.

The end Ce-pair for a desired effect-target depends not only on the pharmaco properties of the agents but also on the maximum infusion rates and concentrations. As shown in fig. 4C, the optimal end Ce-pairs and reaching times to achieve the 90% isobole vary depending on the maximum infusion rates, among other factors.

Once a given target is achieved, different actions can be performed within the DRI model space, Fig. 4D. Simply redefining *x*(*t*0) as the current state of concentrations and establishing a new terminal target vector state ϒ(*t*_*f*_). Specifically, if the current state is desired to remain constant, such as during a maintenance period, the corresponding controls are obtained by setting the new terminal vector state, ϒ(*t*_*f*_), equal to the current state, *x*(*t*0). Fig. 4C, shows the iTCI control for the minimum time to reach the 90% isobole, followed by 10 min of constant Ce concentrations or constant effect. It is worth mentioning that for the particular problem of steady control, the solution provided by iTCI is equivalent to TCI, as the interaction is constant in steady conditions.

### 4.3 Transitions along isoboles

Besides establishing the optimal controls by setting initial and terminal states (*x*(*t*0) and ϒ(*t*_*f*_)), the iTCI algorithm offers the possibility to restrict the control along a viable path (*Γ*) of clinical interest in the Ce-space. Specifically, the iTCI provides control strategies to change between two Ce-pair concentrations belonging to the same isobole while preserving the effect level at every moment.

Fig. 5B illustrates a propofol-remifentanil induction to the 90% isobole followed by a transition along the same isobole. Transitioning along an isobole can be beneficial for adjusting the balance of anesthetics while preserving patient’s state. The minimum time induction to reach an given isobole lead to a triad, (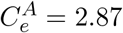 *μ*g ml^*−*1^, 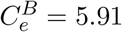 ng ml^*−*1^, *E*_*AB*_ = 90%), where the opioid/hypnotic ratio might be considered relatively large in order to maintain its constancy. In such a situation, for instance, the practitioner could prefer a low opioid/hypnotic target ratio while preserving current patient’s condition. Such control is performed by incorporating the parametric description of the isobole into the iTCI statement, constraint **h**(**x**(*t*), **r**(*t*), *t*) = 0 of eq. (5). The administrations along the isobole requires only a single target concentration since the other drug target is fixed, as both concentrations belong to the isobole. End target along the isobole was set to 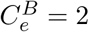ng ml^*−*1^. Below, Fig. 5C shows the equivalent control strategy with lowered infusion rates to 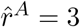 *μ*g min^*−*1^ and 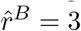 ng min^*−*1^, where the iTCI adapts the infusions accordingly to the new rate restrictions.

**Figure 5:**
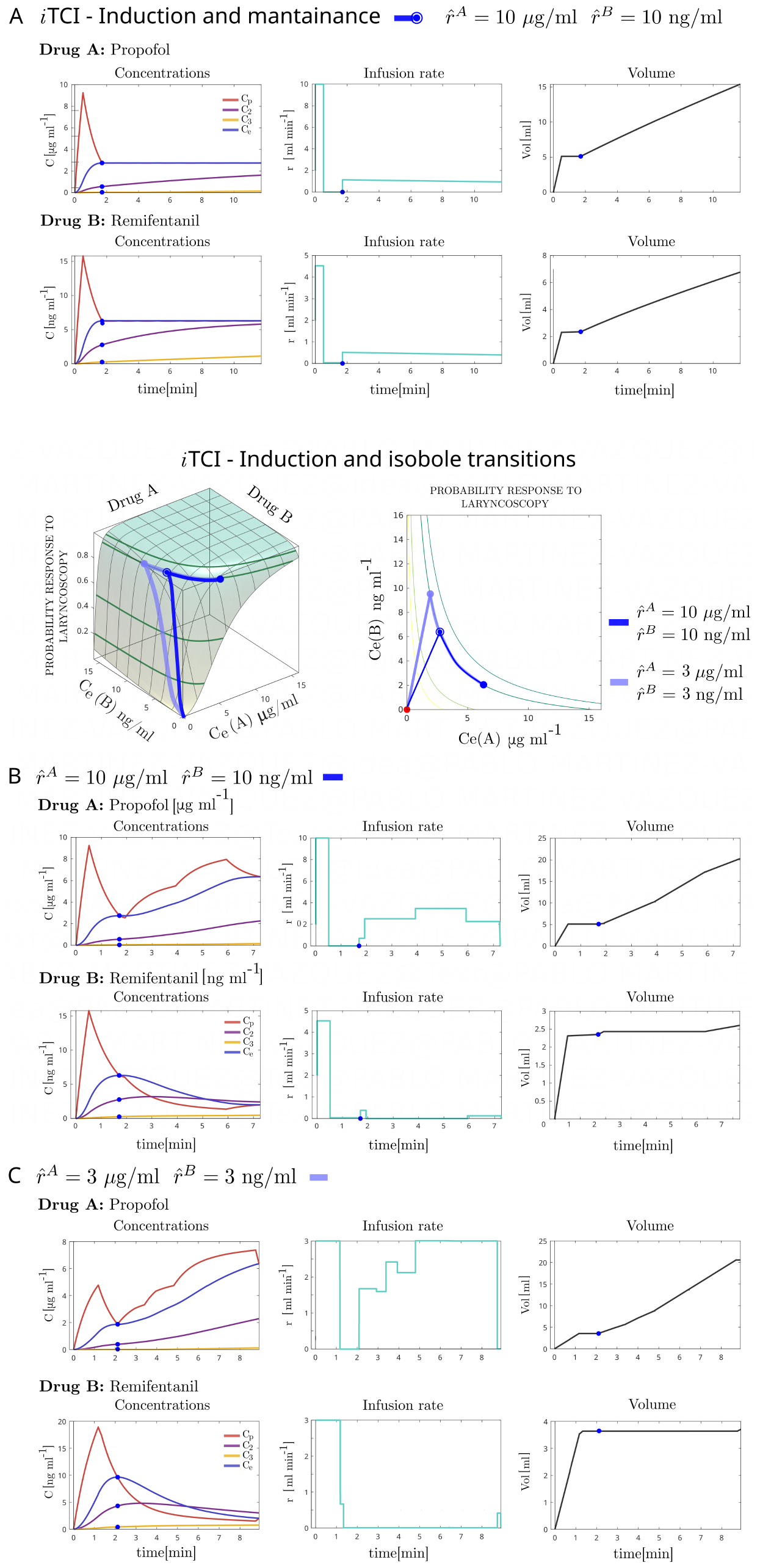
(A) iTCI solution of a minimum time induction to a 90% isobole followed by 10 min stable Ce and Cp target concentrations. (B) Minimum time induction to a 90% target-isobole followed by a transition along the isobole. Once isobole was reached, the end target was set to 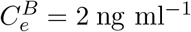 on the 90% isobole. This corresponds to change in the opioid/hypnotic administered ratio while maintaining a common effect. The concentrations, infusion rates, and administered volumes are depicted over time. Maximum infusion rates set to 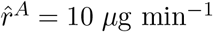 and 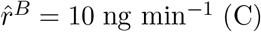 (C) Minimum time induction to a 90% target-isobole followed by a transition along the isobole (target set to 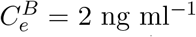 on 90% isobole), with maximum infusion rates lowered to 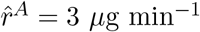 and 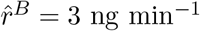.

The iTCI is not limited to isobole restrictions other feasible paths might be clinically interesting such as inter-isoboles path-controlled transitions. However, not all paths in the Ce concentration space are viable. A path that is feasible should comply with the agents’ PK/PD properties and the imposed limitations. The isobole restricted transitions (or other viable paths) cannot be fulfilled with TCI. With TCI systems or manual infusion methods deviations from the isobole (path) are inevitably resulting in undesirable overdosings or underdosings, regarding the targeted effect.

### 4.4 Constraints on the concentrations

Incorporating restrictions on the state variables, *x*(*t*), that describe the agents’ concentrations, can enhance the potential of iTCI. A clinically relevant aspect is the limitation of maximum plasmatic levels, which can help reduce, for instance, undesirable hemodynamics.

During previous induction and transition phases on a given isobole, the plasmatic concentrations of both anesthetics reached high peak values, fig. 5B. Notably, the remifentanil concentration exceeded 15 ng ml^*−*1^ even though the infusion rate was not at its maximum limit. Limiting only the infusion rates is insufficient for efficiently controlling the maximum concentration values. Translating between rates and concentrations is not trivial without the aid of simulations.

Previous infusion and transition on the 90% isobole were re-adapted by adding maximum admissible Cp levels for propofol and remifentanil, 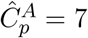*μ*g ml^*−*1^ and 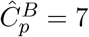 ng ml^*−*1^), respectively, see fig. 6.

**Figure 6:**
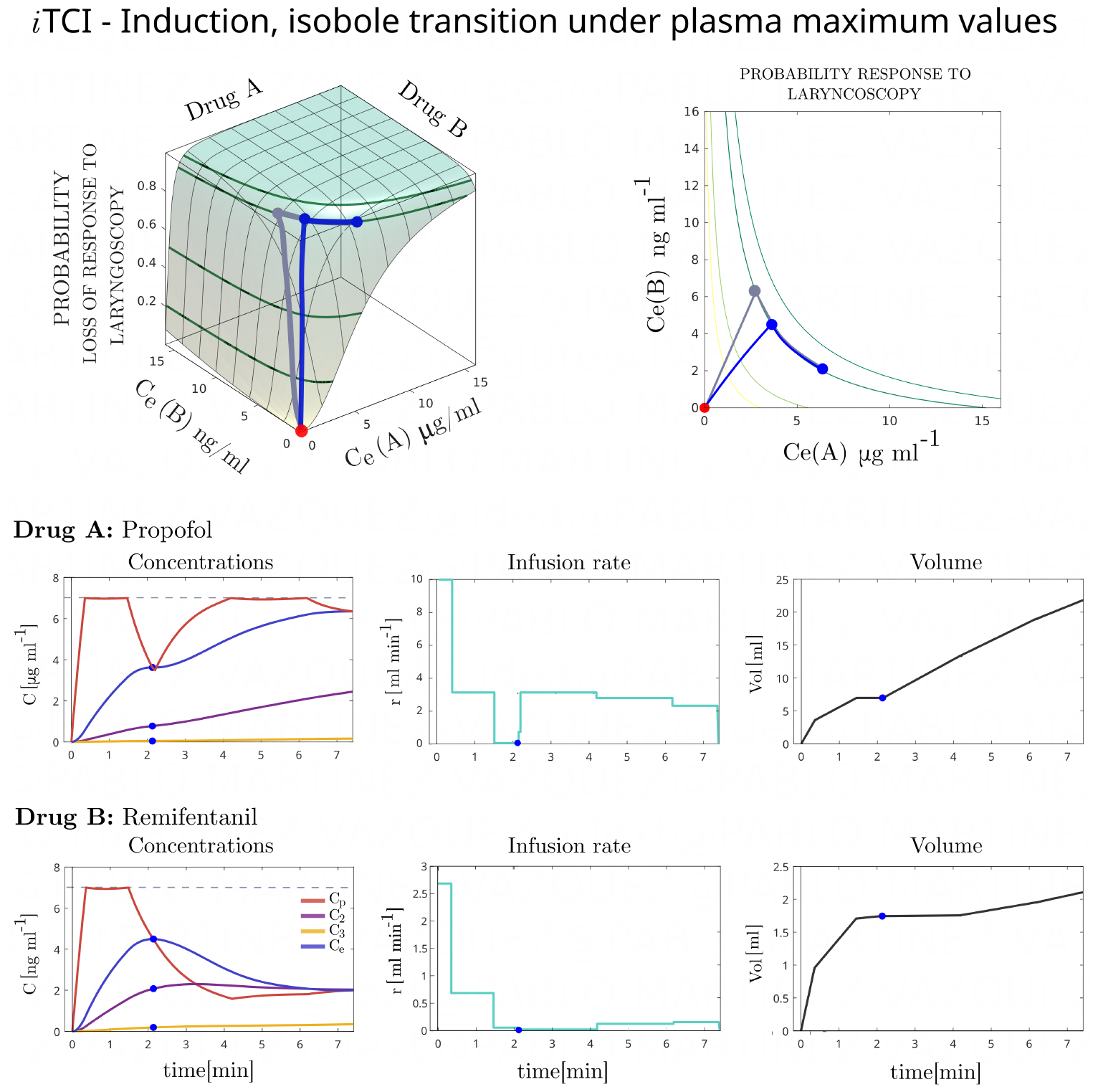
Minimum time induction to a 90% target-isobole followed by an iso-effect transition (end target of 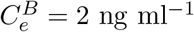 on 90% isobole) with maximum allowed Cp levels. The maximum plasma concentrations, for propofol and remifentanil, were set to 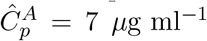 and 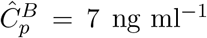, respectively. Concentrations, infusion rares and administered volumens are shown over time. Maximum infusion rates were kept at 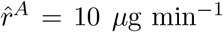 and 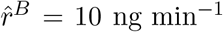. Comparison of this plasmaconstrained iTCI control with the from uncontrained plasma in fig. 5B (gray line), in the DRI model.

Using direct constraints on drug concentrations is safer and more effective than limiting the rate of drug administration. This applies to both multiple and single drug administrations.

### 4.5 Single-drug multi-effect control

Previous examples dealt with controlling two interacting agents, each having different PK/PD properties. However, recent studies^9,31^ pointed out the need to consider different effects with their dynamics even for single drugs. Particularly, for opioids, Abad et al. described that current remifentanil PD models, based on Electroencephalogram (EEG), might reflect more the sedative aspects than the analgesic ones.^9^ They propose extending current opioid PK/PD models to multi-effect multi-effect pharmacokinetic-dynamic (MEPD) models, which are PK/PD models with more than one pharmacodynamic description, see Fig. 7A.

**Figure 7:**
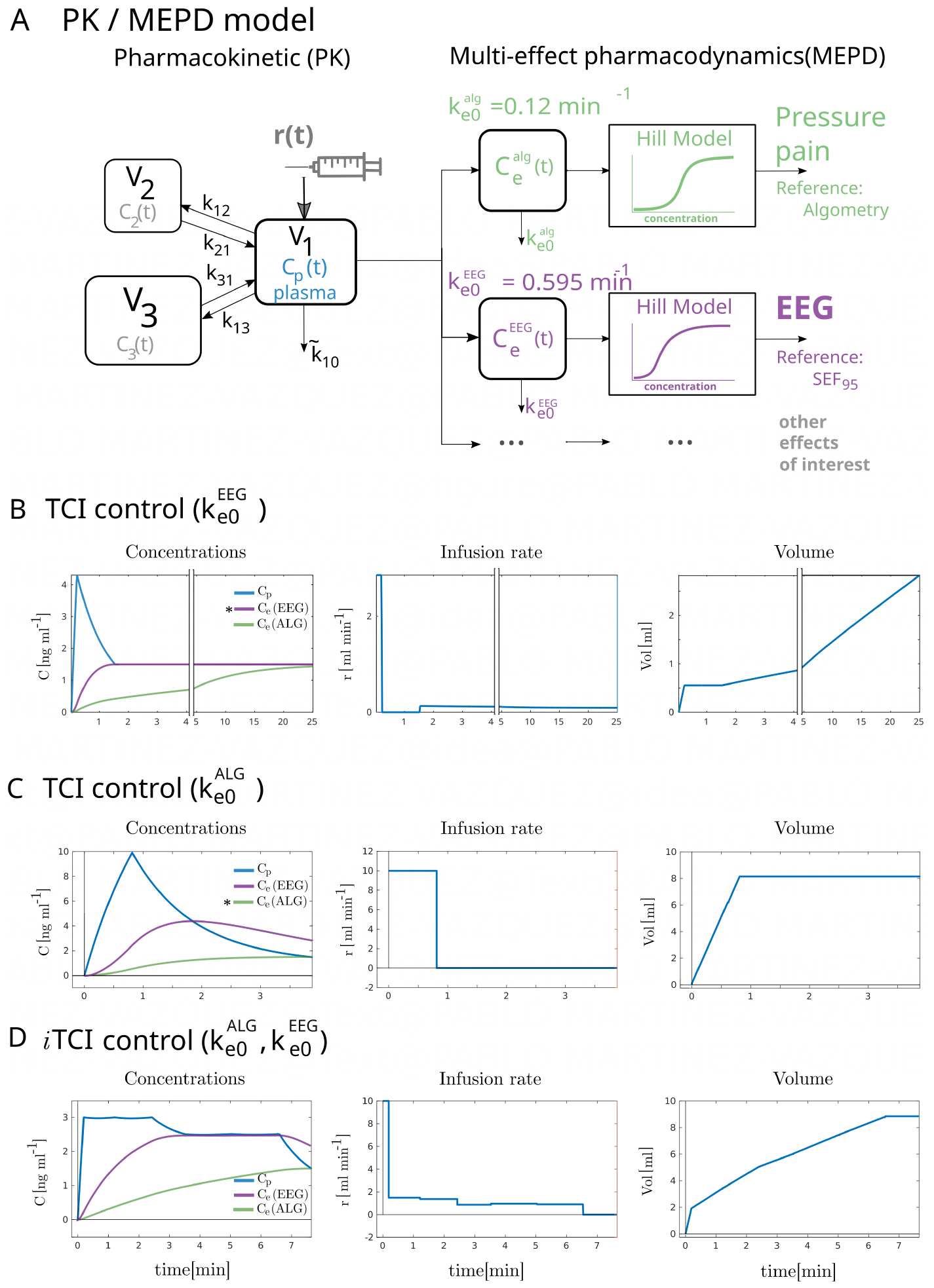
(A) The PK/MEDP model for involves distinct pharmacodynamic models. This example deliniate the effect-site concentration in the brain, estimated through EEG-based measurements, and an effect-site concentration associated with the impact of remifentanil on pressure pain perception. Both effects governed by different time ke0 constants, 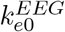 and 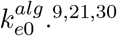 (B) The administration of remifentanil through TCI at a low sedation level, with a targeted effect-site concentration (Ce) set at 1.5 ng ml^*−*1^ and a ke0 constant equal to the EEG-based ke0 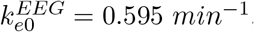, leads to the rapid attainment of the targeted brain-related Ce. However, the measured algometry Ce or the desired level of targeted analgesia is not achieved until at least 20 minutes after the initiation of the infusion. (C) While this results in a faster achievement of the targeted pressure pain analgesia level, the concentration in the plasma (Cp) and the effect-site concentration in the brain (Ce) significantly surpass desirable levels for sedation. (D) iTCI administration of remifentanil at sedation level with pain Ce-target=1.5 ng ml^*−*1^ and maximum concentration levels of 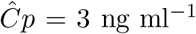 ng ml^*−*1^ and 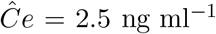 ng ml^*−*1^. The iTCI control is effective in reducing the time required to reach the target analgesic level to approximately one-third of the time needed with the standard TCI method based on the EEG-derived ke0. Additionally, this approach yields to lower peak plasma concentration values.

Abad et al. proposed that at sedation levels, remifentanil infusions would be better characterized by a MEPD model that incorporates the traditional EEG-based PD model estimated by Minto with a 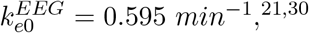 complemented, for instance, with the PDs of remifentanil estimated from pain perception measured by algometry – a better descriptor of the analgesic effect. An effect ruled by a much slower dynamic,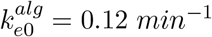. This significant disparity in timings yields in an inconvenient trade-off regarding which effect needs to be targeted, a problem TCI cannot handle adequately^9^.

For example, the administration of remifentanil at sedative levels, (Ce-target=1.5 ng ml^*−*1^), using the standard 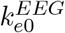 estimation currently in most TCI systems will result in a fast Ce stabilization in brain but in a very long analgesia achievement (approx. 20 min) on patient’s pain perception, Fig. 7B. Alternatively, one might be then tempted to reach more rapidly the analgesic effect by using the algometry-based estimation, 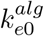, in the TCI. Such a TCI control yield in an administration profile reaching earlier the analgesic target, (approx. 4 min), with the trade-off of excessive maximum Cp and Ce(brain) concentrations for sedative procedures, see Fig. 7C. Both TCI-based options are inconvenient. One requires a long time to reach the desired analgesia and the other being too fast with unfavorable overdoses for a sedation, see Abad et al. for detailed discussion.^9^

The iTCI offers a better administration. Including the two two *k*_*e*0_s in the iTCI statement, significantly shorter analgesia target time is achieved while limiting the maximum admissible drug concentrations. Fig. 7D depicts the iTCI control based on algometry dynamics while limiting the maximum concentration levels of Cp and Ce in brain to 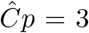 ng ml^*−*1^ and 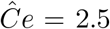 ng ml^*−*1^, respectively. Limits more suitable for sedations that the ones obtained in the TCI based on the 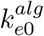, Fig. 7C. The iTCI allows to reduce largely the time to reach the analgesic effect while keeping admissible concentration levels. This example demonstrates the capability of the iTCI framework over standard TCI, even in single drug controls. It can integrate multiple drug effects in the control problem by directly limit the maximum concentration levels.

The asterisk (*) denotes the selected *k*_*e*0_ applied in each TCI control. Maximum infusion rate was set to 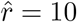 ng min^*−*1^.

## 5 CONCLUSIONS

The iTCI framework is a novel method for administering multiple anesthetics that interact with one another. It involves combining the pharmacokinetic and pharmacodynamic models of the coadministered drugs and their interactions, expressed by DRI models, into a single optimal control problem. This approach enables concurrent optimal infusions, which can be defined based on target goals, either as concentrations or effects.

The iTCI allows controlled infusion strategies of clinical interest that the single-drug TCI systems cannot perform. Examples of iTCI capabilities were presented for two of the most used agents in GA, propofol and remifentanil, interacting both synergistically.

iTCI is a control strategy that assumes all desired Ce-targets are integrated into a single target state. This allows iTCI to achieve the desired Ce-targets simultaneously, reducing the risk of unnecessary premature partial target achievements. It achieves this by infusing faster-acting drugs at a slower rate than TCI, which helps to adapt to the slower-acting agents and reduce their overall administered volumes and maximum plasma concentration levels.

In iTCI, the desired targets can be defined in terms of effects. Incorporating effects into the control strategy allows, for example, to perform the fastest infusions to a target isobole. Additionally, the infusions can be constrained to follow specific paths within the Ce-space. The restriction of changes between two iso-effect administration regimes along its isobole is clinically relevant when practitioners want to change between drug delivery regimes while preserving the effect. Such control is not possible in TCI, leading to overdosing or underdosing from undesirable deviations of the target isobole.

Constraints on the maximum infusion rates can be set as in TCI, but additional constraints can be applied directly to state variables, such as plasma concentrations. Direct constraints on plasma levels provide safer and more accessible control management than limiting only the infusion rates since the impact on concentration levels from rate limits is not trivial.

The iTCI can function as a TCI in single-drug administrations. However, single-drug iTCI infusions can include different pharmacodynamics related to various drug effects. This may be relevant for opioids. Recent research conducted during sedation has demonstrated pharmacodynamic discrepancies between the analgesic effects of remifentanil and its impact on brain activity. This suggests that PK/PD models should be expanded to incorporate diverse pharmacodynamics of interest. TCI is unable to control infusions under several exerted pharmacodynamics.

The iTCI requires PK/PD models and interaction descriptions as minimum requirements. Therefore, iTCI is not restricted to IV agents only. Gases such as sevoflurane with a PK/PD model^32^ could be controlled along with other anesthetics like remifentanil, which interact synergistically.^33^

The iTCI is not limited to pharmacokinetic linear models, 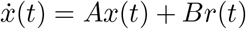, it can operate on general dynamic pharmacometrics models, 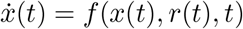, that might enhanced the accuracy of drug dynamics estimation.

In summary, the iTCI methodology presents a single, all-encompassing nonlinear dynamic optimization control problem. This allows for safer and more flexible optimized concurrent administration profiles than traditional TCI solutions. This opens up new possibilities for future multi-anesthetics controlled administrations where aspects such as synergies can be taken into consideration for better anesthesia.

## 6 APPENDIX SUPPLEMENTARY MATERIAL

### A-1 PD/PK Schnider and Minto models

**Table A-1:**
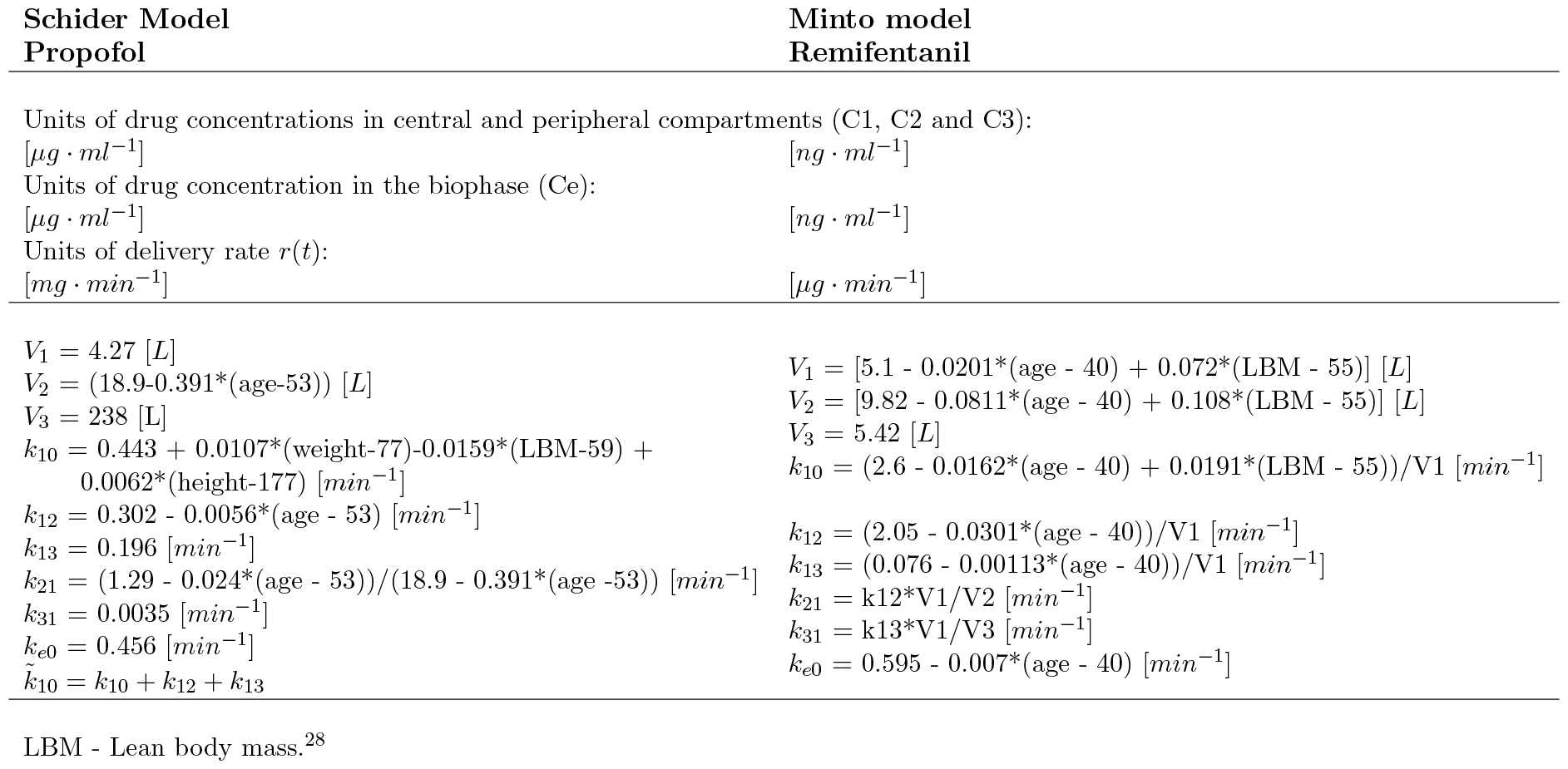
Propofol and remifentanil PK/PD parameters’ estimates by Schnider and Minto, respectively. ^29,30^

### A-2 iTCI algorithm description

The iTCI solution relies on the finite element method (FEM) method. The FEM is a systematic approach to approximating continuous functions that breaks down a problem’s domain into a finite number of points and subdomains. These subdomains (finite elements) are represented by simpler functions (e.g., polynomials) held at specific points called nodes, which are connected at their boundaries. They hold piecewise and local approximations of the function, which are uniquely defined in terms of values held at their nodes. The solution to the iTCI statement, eq. (5), consists of the assembled elements that fulfill the boundary continuity conditions between adjacent finite elements and all requested conditions. Thus, the PK/PD and interaction models are met, along with the initial and terminal constraints and any added limitation on controls (infusion rates) and states (concentrations and effects).

Controls and states are discretised, within the interval *t ∈* [*t*_0_, *t*_*f*_], by i = 1, *…*, NE finite elements, fig. 2. Each i-th finite element approximate the control and states solutions, on the interval *t ∈* [*ξ*_*i*_, *ξ*_*i*+1_], with Lagrange polynomials of order K+1 and K, respectively, eqs. (A-1a) and (A-1b). The difference in orders is due to the existence of initial conditions for the states in each of the finite elements.

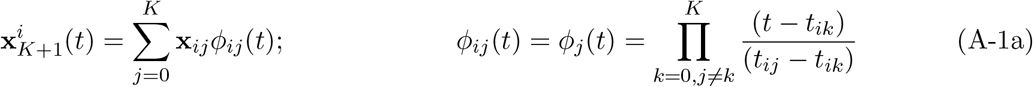

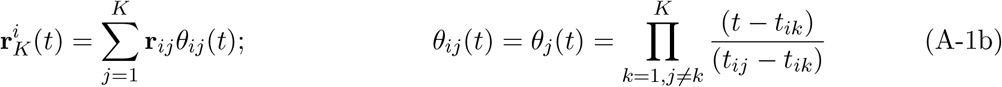

The orthogonality of Lagrange basis, *ϕ*_k_(t_j_) = *δ*_kj_ (*δ*_*kj*_ stands for the Kronecker delta), leads to the property 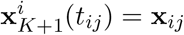, which allows for a direct bounding of the states and controls. Particularly, allowing the definition of path constraints on the problem formulation.

**Figure A-1:**
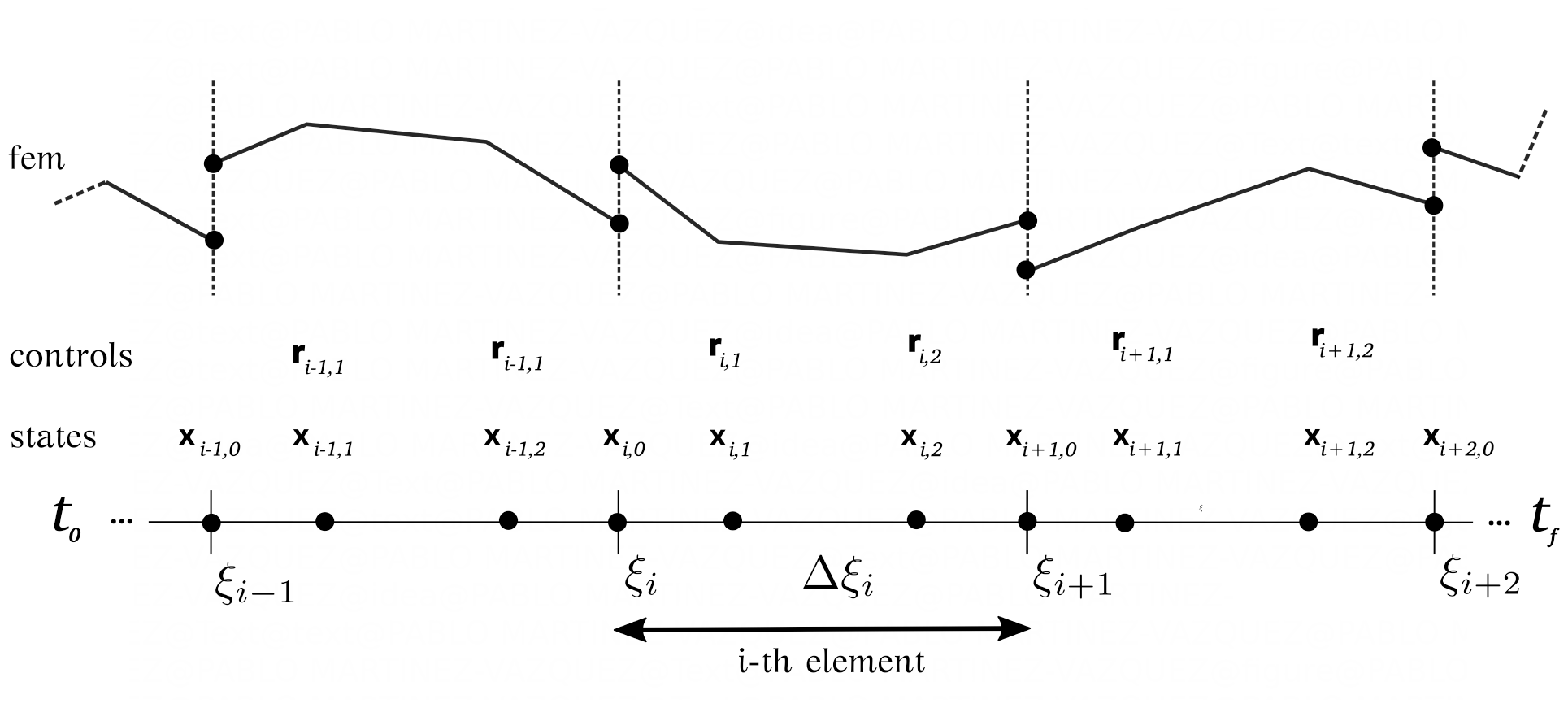
FEM are continuous functions that breaks down a problem’s domain into a finite number of points and subdomains. Controls and states are discretised. Finite-element collocation.

The collocation of the knots, *ξ*_*i*_ and *ξ*_*i*+1_, bounding the i-th finite element of length Δ*ξ*_*i*_ = *ξ*_*i*+1_ *− ξ*_*i*_ and times *t*_*ij*_ within the interval *t ∈* [*t*_0_, *t*_*f*_], are derived from the roots of Legendre polynomials mapping each finite element in the unit interval, Δ*ξ*_*i*_(*τ ∈* [0, 1]). Such a distribution of the NE finite elements preserves orthogonality properties leading to an efficient and accurate approximation of the controls and states profiles.^23,24,26^

Continuity across adjacent finite elements is imposed with eq. (A-2a) on the states, or equivalently eq. (A-2b). However, continuity is not applied on the controls to allow discontinuities at the finite elements endpoints.

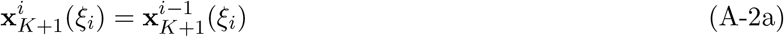

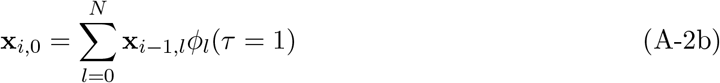

Substitution of eqs. (A-1a) and (A-1b) into the dynamical system 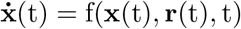 leads to the approximation residuals, R(t_ij_), at the discretization points t_ij_ = *ξ*_i_ + Δ*ξ*_i_*τ*_j_ of dynamic system, where 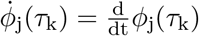. Residuals must be equal to zero, section A-2.

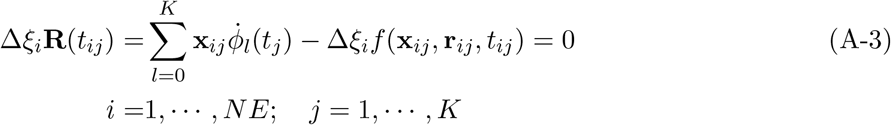

The dynamic nonlinear optimization problem with constraints, eq. (5), when discretized on finite elements that fulfill states continuity conditions and bounds on both profiles at the knots becomes the algebraic NLP, eq. (A-4):

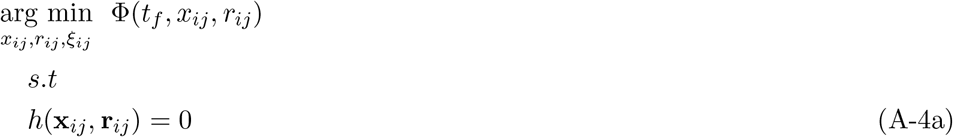

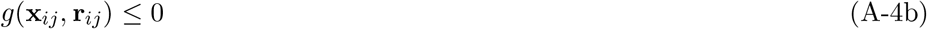

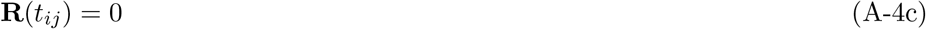

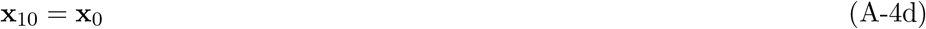

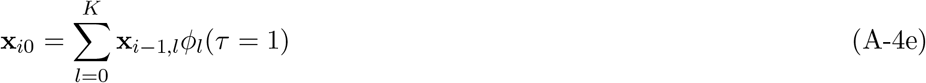

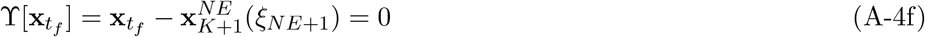

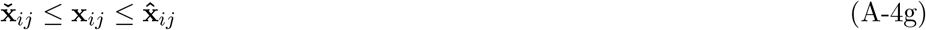

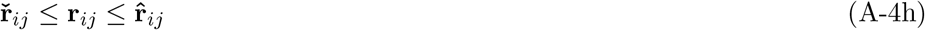

where *i* = 1, …, *NE, j* = 0, …, *K*, and 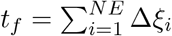. Basis functions, *ϕ*_*j*_(*τ*), and their derivatives, 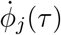, in eq. (A-2b) and section A-2 or eq. (A-4c), are calculated beforehand, since they depend only in the Legendre roots locations. The NLP can be solved numerically with a general-purpose SQP solver,^27^ which provides the iTCI solution.

## Glossary

Ce: effect-site concentration.
Cp: plasma concentration.
DRI: dose-response interaction.
EEG: Electroencephalogram.
FEM: finite element method.
GA: general anaesthesia.
iTCI: interaction target-controlled infusion.
IV: intravenous.
LOR: loss of response.
MEPD: multi-effect pharmacokinetic-dynamic.
NLP: non-linear programming.
PK/PD: pharmacokinetic/pharmacodynamic.
SQP: successive quadratic programming.
TCI: target-controlled infusion.

